# How the characteristics of a virtual environment affects the perception of travel distance through it

**DOI:** 10.1101/2025.09.10.675295

**Authors:** Ambika Bansal, Meaghan McManus, Laurence R. Harris

**Affiliations:** Centre for Vision Research, York University

## Abstract

Although simulated self-motion through virtual environments has been widely used to investigate perceptual odometry, the characteristics of the virtual environments used, and the reported results have varied greatly. Here, we systematically vary the characteristics of the environment through which observers are moved in order to explore the effect of (1) the structure of an environment including the presence and texture of a ground surface, (2) the naturalism and scale of an environment, (3) colour, and (4) the density of a starfield and how it might affect perceived travel distance. In all four experiments, participants were visually moved forwards through a virtual environment and perceived travel distance was estimated by either (1) stopping at the location of a previously seen target (the Move-To-Target Task) or (2) adjusting the position of a target to indicate a previously travelled distance (the Adjust-Target Task). Data were analyzed in terms of gain (perceived travel distance/actual travel distance). Results show no significant differences that depended on the structure of an environment or on the presence or absence of a ground surface (Experiment 1), or on the naturalism of the environment (Experiment 2), or on whether the environment was in colour or in black and white (Experiment 3). However, there was a small effect of the texture of the ground surface and of the scale of the environment. In Experiment 4, we show that there may be a very low ceiling effect in the density of a starfield needed to accurately estimate travel distance. Together these experiments have implications for the design of real and virtual environments where perceived motion is important and will enable us to further predict our perception of moving through an environment.

## Introduction

Understanding how we interpret our environment is crucial to fully appreciate how we perceive our movement through it. Although visually induced self-motion has been widely used to investigate the perception of travel distance (how far a person perceives themselves to have traveled through an environment), the visual characteristics of the virtual environments used vary greatly between studies. There are various parameters of a virtual environment (structure, texture, ground surface, colour, luminance, distance to walls, etc.) that have been shown to contribute to the processing of optic flow (Bubka & Bonato, 2010; Seno et al., 2010; Tamada & Seno, 2015; Vaziri-Pashkam & Cavanagh, 2008). Although these variables have been shown to clearly modulate self-motion perception and therefore the perception of travel distance, these parameters are rarely taken into consideration when designing self-motion experiments or virtual gaming environments. Understanding how these various visual field characteristics might affect the perception of travel distance is a crucial element in improving our understanding of this system.

Optic flow alone can induce an illusionary sensation of self-motion (Fischer & Kornmiller, 1930; Brandt et al., 1973). A real-world scenario in which it is common to experience vection is when sitting in a stationary train while visually watching another train move past you through the window. You will get the illusion that you are moving in the opposite direction of the passing train even though your train is stationary. This illusion of self-motion is known as vection and has been used in many studies as a correlate of self-motion perception. However, one can still experience self-motion and perceive one’s travel distance without experiencing vection (i.e., without the sensation that one is actually moving). Using a similar environment to our virtual corridor in Experiment 1, McManus and Fiehler (2025) have shown that when participants feel vection, the felt speed of self-motion and perceived travel distance are correlated, and if no vection is experienced then the two are not found to be correlated. Although it should be noted that in majority of their trials, participants did not actually experience any vection. Many studies investigating how the characteristics of an environment affect self-motion perception have used the self-reported magnitude of the sensation of vection as a measurement, but here we are specifically concerned with the effect of the environment on perceived travel distance.

Humans have evolved to move around on a stable ground surface and are tuned to pick up relevant information about their self-motion largely from the ground (Gibson, 1979). Understanding how we process the movement information provided by the ground surface is crucial not only to understand our naturalistic movements, but also when considering the processing involved when experiencing modern forms of motion such as in driving or aviation and the safety issues involved in these unnatural forms of self-motion. Whether it is taking off or landing an aircraft, driving or walking, optic flow received from the ground surface is an important feature in the perception of self-motion. Bian and colleagues (2005) introduced the idea of a “ground dominance effect” in the perception of a 3-D scene. When comparing information across all four environmental surfaces (ground, ceiling and the sidewalls), participants are consistently dominated by visual information from the ground surface. Previous research has found that projecting optic flow patterns onto a ground surface results in body sway corresponding to the flow direction (Flückiger & Baumberger, 1988). This is true for both children and adults (Baumberger et al., 2004). A previous study has manipulated the size, position, and speed of the components of a random dot stimulus on a floor projection to investigate the effects of these properties on linear vection and body sway (Tamada & Seno, 2015). Results show that some of the stimulus properties of the ground surface had an effect on both vection and body sway. Trutoiu and colleagues (2009) tested the effects of a floor projection on the perception of vection for linear forward, linear backward, circular left, and circular right directions. They found that adding a floor projection of a starfield significantly increased the sensation of linear vection, but not circular vection. This ground dominance effect has also been replicated using visual search tasks (McCarley & He, 2000). These observations provide further evidence that visual-field characteristics can modulate optic flow processing. Surface texture has also been shown to influence vection. More specifically, surface textures containing high spatial frequencies with realistic lighting (e.g., bark, wood) enhance vection, whereas smooth or reflective surfaces (e.g., glass, metal) provide less motion-relevant visual information, and therefore tend to reduce vection strength (Morimoto et al., 2019). Although the ground surface and its texture seem to be critical features in visual perception, little has been done to examine its effects on perceived travel distance. In Experiment 1, we test this question directly. Previous research from our lab has found variations in perceived travel distance in which participants felt they had moved further when emersed in a virtual corridor (Bansal et al., 2024) compared to studies using less structured environments (e.g., Bury et al., 2020; McManus & Harris, 2021). These “less structured environments” included a starfield stimulus and a “lollipop field”. Although it is difficult to fully quantify environmental structure, the star field and lollipop environments are considered to be less structured because in the starfield there were no polarized cues to orientation, and in both environments, the objects were arranged with no systematic structure and can thus be considered more abstract and less naturalistic than, for example, a realistic virtual simulation of a corridor. More detailed environments have also been shown to lead to more accurate walking trajectories than less detailed environments (Wood et al., 2000). Though differences in travel distance estimates have been observed across studies, no one has yet systemically tested the effects of environmental structure on perceived travel distance.

Extracting the optic flow created by moving in natural environments is complex because of the continuously changing distance to all the objects in the environment producing continuously changing angular velocities of the images of all the objects in the field. One possible confounding variable for studies investigating environmental structure and the ground surface on vection is simply the ability of participants to extract an appropriate amount of optic flow information. In more complex environments, it could be that denser optic flow information could lead to a stronger sensation of vection, and a further perceived distance of travel. In Experiment 4, we directly investigate the starfield density needed to accurately estimate travel distance.

More naturalistic environments have been shown to lead to a stronger magnitude of vection. Visually moving through a photo-realistic three-dimensional simulation of a town presented in a large-screen virtual environment induces a stronger sensation of vection compared to moving through a star field (Trutoiu et al., 2009) or through abstract stimuli (Riecke et al., 2005; 2006). These researchers describe how the “believability” of a visual scene or environment may contribute to one’s sense of presence, and therefore the strength of vection. Some researchers have also tested how the semantic meaning of a scene may affect vection (Ogawa & Seno, 2014; Seno & Fukuda, 2012). One such study used the train illusion described above to induce vection, where participants viewed animated scenes of a train passing (Seno & Fukuda, 2012). The train-context triggered faster onset and longer vection duration compared to abstract stimuli moving passed at the same speed. Another study found that when perceiving a pattern of optic flow in the downward direction there was a reduction in vection when the falling objects were recognizable compared to meaningless, abstract dots, despite identical visual motion patterns (Ogawa & Seno, 2014). In a second experiment from the same study, vection was also reduced when participants were holding an umbrella (feeling sheltered – while viewing these falling objects) compared to when they were holding a sword (with no contextual meaning), again, despite identical motion. These studies provide evidence that vection is not solely driven by bottom-up visual motion. It is clearly also modulated by top-down factors like semantic context and scene interpretation. Here, we use variations in “naturalism” to manipulate semantic meaning. However, it is difficult to quantify “naturalism”. In this study, we describe a scene as natural if it obeys the rules of the natural world (i.e., do the scale and orientation of the scene’s objects align with the real world?). While the relationship between vection and self-motion is not well researched, it is possible that the enhancement to vection by more naturalistic environments (as described above) might also lead to changes in perceived distance traveled during a self-motion task. No one has yet investigated how the naturalism of a scene may affect the perception of travel distance. In Experiment 2, we test this question directly.

Colour has also played an increasingly recognized role in self-motion perception. Similar to the work on naturalism and vection perception, past research has found that more “complex” environments lead to improved sensation vection where complexity was increased by increasing the visual-field characteristics of the drum lining of an optokinetic drum, for example, by changing a stripe pattern to a checkerboard pattern or by adding different colours to the stripes (Bonato & Bubka, 2006). One study tested how chromatic colour, specifically red versus green, affected vection over a series of seven experiments (Seno et al., 2010). They found that: (1) a red background consistently produced weaker vection compared to a green background, (2) red moving dots on black also yielded reduced vection compared to green dots on black, (3) a moving red grating induced weaker vection than a green grating, (4) combining red dots on red background led to even weaker vection, while green-on-green produced stronger vection, and (5) these effects were not due to luminance artifacts. Together, this series of experiments suggest that red visual stimuli suppress vection, and that colour does have an effect on self-motion perception. In a carefully luminance-controlled environment, chromatic dots (i.e., dots with colour) produce stronger vection compared to achromatic dots (i.e., gray and white dots), (Seya et al., 2015). Another study showed that coherent colour modulation, irrespective of colour type, results in significantly reduced vection strength, meaning longer onset latencies, shorter durations, and lower magnitude scores (Nakamura et al., 2010). These studies clearly support the idea that the colour of an environment can affect self-motion perception. Therefore, in Experiment 3 we examine how the colour of an environment may affect the perception of travel distance.

Many studies on the perception of travel distance have used either the Move-To-Target (where participants estimate travel distance by stopping at the location of a previously seen target) or the Adjust-Target task (where participants adjust a target to indicate the distance of a previous movement). Few have used both to measure travel distance estimates. Since we have previously found differences in travel distance estimates between these two tasks (Bansal et al., 2024), we used both tasks here to measure travel distance estimates in the present study.

The overarching objective of the present study was to investigate how different characteristics of a virtual environment may affect the perception of travel distance through it. In Experiment 1, the objective was to test the effects of structure, the presence of a ground surface, and its texture on perceived travel distance in virtual reality. In Experiment 2, the objective was to test whether there was an effect of naturalism on perceived travel distance. In Experiment 3, the objective was to test how colour affects the perception of travel distance. In Experiment 4, the objective was to test how the density of a starfield can affect perceived travel distance.

We hypothesize that since the variables of the first three experiments have already been shown to affect vection and self-motion perception (as reviewed above), the presence and texture of the ground surface, the addition of more structure and more naturalism, and the presence of colour would lead to larger estimates of travel distance, meaning participants will feel like they have moved further in these conditions. We also hypothesize that there may be a ceiling effect with the starfield density needed to make accurate travel distance estimates. This study will give us a deeper understanding of how the specific components of visual motion contribute to our perception of self-motion.

## Experiment 1: Environmental Structure, Texture, and the Ground Surface

### Methods

#### Participants

Eighteen subjects (9F, 9M, mean age 19.4 yrs, SD ±2.1) participated in this study. The recruitment period was between January 21st, 2023 and April 3rd, 2023. All participants were recruited using the Kinesiology Undergraduate Participant Pool at York University. The protocols used in this study were approved by the York Human Participants Review Sub-committee (#e2021-407) and conducted in accordance with the Declaration of Helsinki. All participants gave prior informed written consent and were naive to the purpose of the study.

#### Apparatus

The equipment used in this experiment was a virtual reality (VR) head mounted display (HMD) and Alienware computer (Intel Core i7-8700K CPU, 3.70 GHz). The virtual stimuli were presented via an Oculus Rift CV1 (Oculus VR; 90 Hz refresh rate, 1080 × 1200 resolution per eye).

#### Stimuli

The experiment was performed in virtual reality using visually induced self-motion, while the participants remained physically stationary. There were two virtual environments in which this experiment was performed. The first, more structured environment was a virtual corridor that was 1.86m x 1.86m, with the simulated eye height set to the center of the corridor at 0.93m above the ground (Fig 1, top row). The second was a less structured “starfield” environment, with a starfield density of 3,000 stars and a ground surface (Fig 1, bottom row). Each star in the starfield environment was 3.5m in diameter and had an infinite lifetime. In both environments, there were three ground surface conditions. The first had a ground surface with a virtual grass texture, the second had a vertical striped pattern on the ground surface with no texture which therefore did not provide any optic flow information, and the last condition had no ground surface at all. In the no-ground-surface condition, the block started by having the ground “drop” before participants began the first trial to stress how there was no visible ground surface while they were experiencing the simulated movement.

**Fig 1.**
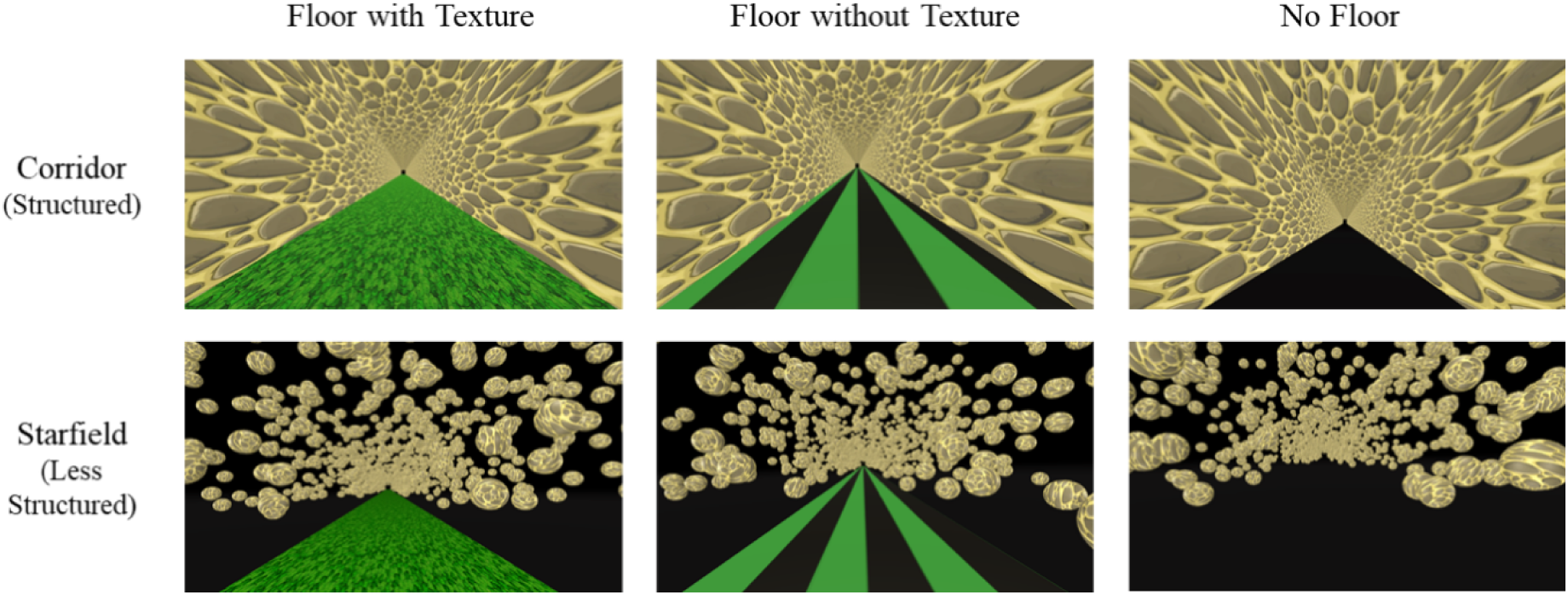
Visual scenes seen inside the virtual reality headset. The top row displays the virtual corridor (more structured) environment. The bottom row displays the starfield (less structured) environment. The first column shows both virtual environments containing the ground surface with a “grass” texture. The second column shows both environments containing the ground surface without a texture (plain green). The last column shows both environments without the ground surface.

There were six different conditions (2 environments x 3 ground surface options). In the starfield environment (bottom row of Fig 1.), the stars were only present above the horizon in the visual display, and had the same texture as the virtual corridor, to more closely match the visual display characteristics of the virtual corridor. The starfield environment was arranged such that the stars could not hit the participant. As to not create a “box” of stars around the participant, the stars were placed at a random location to the left, right, or above them. On the left, the stars were placed anywhere between −25m and less than a random number between −4 or −6m, on the right, the stars were placed between +25m and greater than a random number between +4 and 6m, and on top, the stars were places between +25m and greater than a random number between +9 and +11m. The stoney texture on the walls of the corridor was arranged such that the stones were randomly placed in a non-repeating pattern. Participants were visually moved at a constant velocity of 5 m/s, which was the velocity used in our previous study (Bansal et al., 2024). The visual stimuli were always presented stereoscopically and world-fixed, such that the visual environment updated with the participant’s head movement, though participants were asked to keep their head straight.

#### Move-To-Target Task

Each trial started with a simulated target (a black-and-white checkered pillar measuring 1m x 5m) presented straight in front of them (Fig 2). The target was presented in a separate sparse black environment with a black and white striped ground surface, so participants did not have any landmarks to use as the target’s location during the “movement phase”. They then pressed a button on a mouse which triggered the target to disappear and moved them to the start of the virtual corridor or starfield depending on the condition. Then motion in the virtual corridor or starfield was simulated with optic flow. Participants indicated when they felt they reached the now invisible target by pressing the button once more.

**Fig 2.**
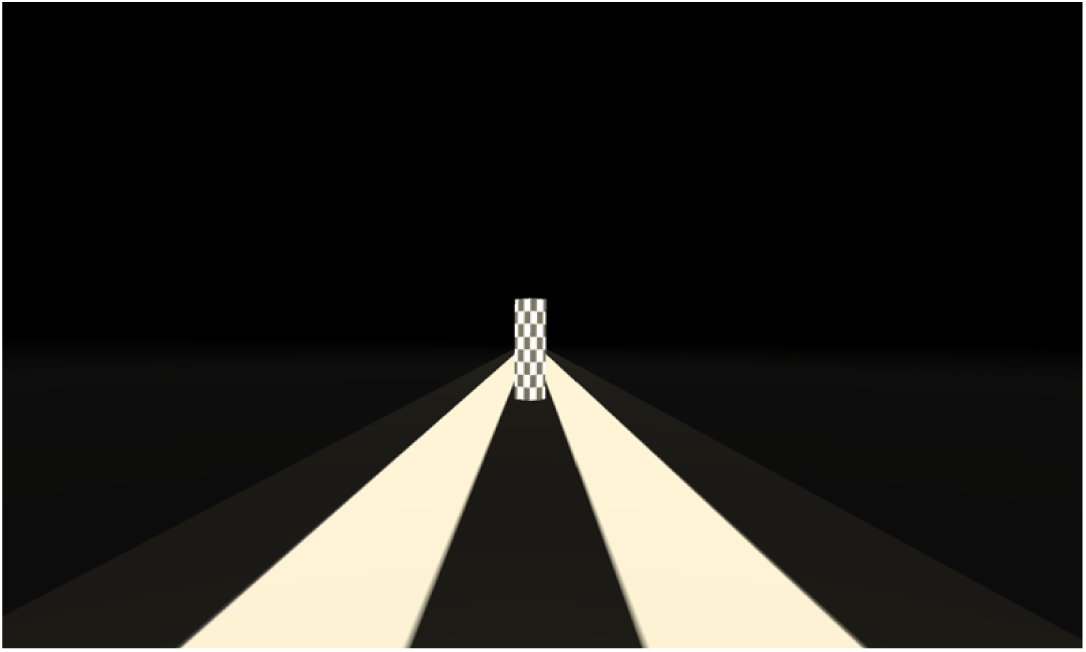
Visual display of the target environment. For the Move-To-Target task, this was the target presented in front of the participants at the beginning of each trial. For the Adjust-Target task, this was the target that participants had to adjust to the distance they felt they had been moved.

#### Adjust-Target Task

Participants sat in the same virtual environments as for the Move-To-Target task (either the starfield or corridor depending on the condition) and began by pressing a button. They then experienced simulated movement through a pre-specified distance. Once they had traveled this distance, they were teleported to the sparse black and white environment with the same target as for the Move-To-Target task. Participants then used the up and down arrow keys of a keyboard to slide the target back and forth along the corridor until it was as far away as the distance through which they felt that they had just been moved. They pressed the space bar to end the trial.

#### Procedure

Participants were first asked to sit in a chair, while the instructions were explained to them by the experimenter. At the beginning of each task, participants were first given a practice session which included 6 trials (2 trials in each of the three ground surface conditions) at randomized distances between 5-40m. Once the practice was completed, the full version of each task was run.

This study was a within-subjects design, such that every participant completed both the Move-To-Target and Adjust-Target tasks. The order in which the tasks were completed was counterbalanced such that half completed the Move-To-Target first and the other half completed the Adjust-Target task first. For each task the participants experienced two structured environments: corridor or starfield, which each had three ground surfaces (grass textured, plain green, or no ground surface). Within each of these conditions the environments moved at 5 m/s through one of twelve target distances (5, 8, 11, 14, 17, 20, 23, 26, 29, 32, 35, 40 m). Each condition was presented only once. All conditions were randomized to cancel out any order effects. The tasks were blocked by surface and environment conditions and the order in which these blocks were presented was also randomized. The instructions were presented in the HMD again at the beginning of each block and at those times the participants were able to take a break if needed. The whole experiment took about 30 minutes (15 minutes for each of the two tasks).

#### Data Analysis

Each participant completed 144 trials (2 structured environments x 3 ground surface conditions x 12 distances x 2 tasks). First, an outlier removal was completed. The outlier removal was performed at the group level for each distance in every condition (2 environments x 3 ground surface conditions x 2 tasks). Any data less than the ‘Lower Quartile - 1.5 x Interquartile Range’ or above the ‘Upper Quartile + 1.5 x Interquartile Range’ were removed. Out of 2,592 data points, 59 data points were removed.

The ratio of travel distance to target distance (gain: output/input) was then calculated for each trial for both tasks. In the Move-To-Target task, this translates to the target distance (how far they thought they had moved) divided by the distance travelled (the amount they were simulated to move) before pressing the button to stop:

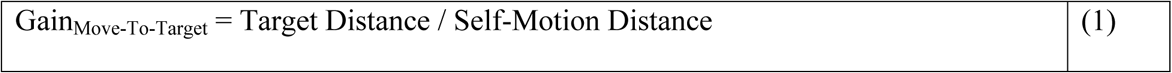

For the Adjust-Target task, this translates to where participants adjusted the target to (how far they thought they had moved) divided by the distance that participants were moved through (the amount they were simulated to move):

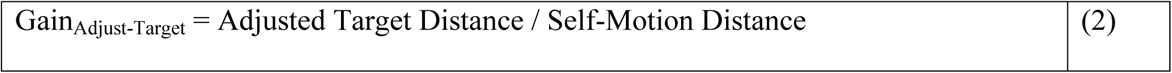

Perfect performance in both cases would be a gain of 1. In both cases, a gain greater than 1 would imply that participants felt like they had moved further than they had, and vice versa. Before testing any differences between conditions, we tested whether any effects for a condition differed between MTT and AT tasks. Since we did find differences between tasks, we analyzed the tasks separately. A Linear Mixed Model was performed independently for both tasks using the lme4 package (Bates et al., 2015) for R (version 4.3.0.) on the gains. To determine the most appropriate model structure, we used Barr et al.’s (2013) model comparison approach. We started with a maximal model including all relevant experimental variables (environment structure, and ground surface condition) as slopes per participant and compared to simpler models until no significant differences were found between models. Random slopes for environment structure and ground surface condition per participant were kept for both models. The fixed effect structure was chosen as a function of our hypotheses, where we were interested in the main effects of environment and ground surface. We did not include an interaction between environment and ground surface because we had no specific hypotheses about the interaction term. The final model structure for the gains LMMs reads as follows:

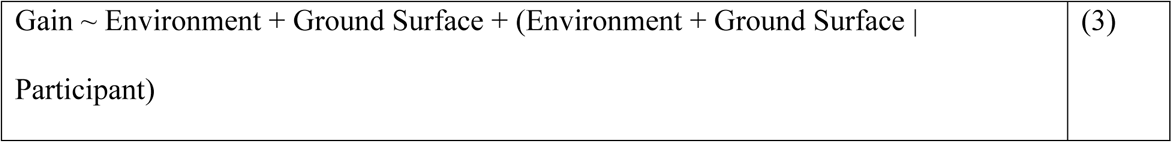

We then computed bootstrapped confidence intervals at an alpha level of 0.05 to test for statistical significance using the confint() function from the base R with the “boot” argument and default settings otherwise. All data and data analysis can be found at https://github.com/ambikabansal/Virtual-Environments.

## Results

### Effect of Environment Structure

#### Move-To-Target Task

The top row of Table 1 shows the means and standard deviations of the gains for the Move-To-Target task. Table 2 shows the results from the linear mixed model for the effect of Environment Structure on the gains of the Move-To-Target task. We found no significant differences between the corridor and starfield environment conditions. The gains are shown in Fig 3.

**Fig 3.**
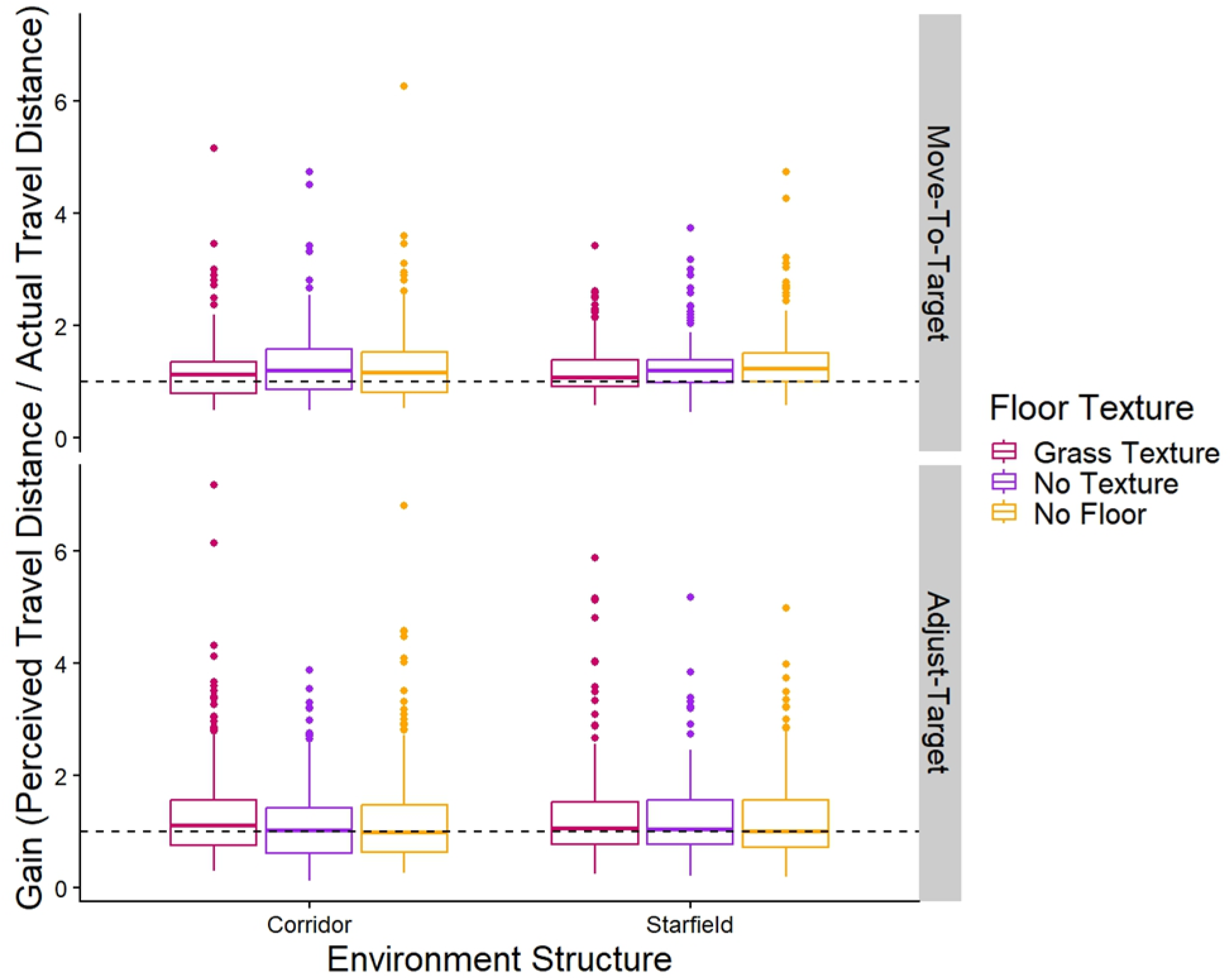
Gains. Box plots of the group gains for both the Move-To-Target (top row) and Adjust-Target (bottom row) tasks for each Environment Structure and Ground Surface Condition. The middle line represents the median, the boxes extend from the first quartile to the third quartile, the whiskers extend up to 1.5 times the interquartile range, and the outliers are shown as individual points beyond the whiskers. The grass texture is represented in maroon (left most), no texture in purple (second from left), and no ground surface in yellow (right most). The black dashed line represents perfect performance (gain of 1).

**Table 1:** Means and standard deviations of the gains for the different Environmental Structure conditions.

|  | Corridor | Starfield |
| --- | --- | --- |
| Move-To-Target Task | 1.24 ± 0.61 | 1.28 ± 0.51 |
| Adjust-Target Task | 1.25 ± 0.88 | 1.26 ± 0.77 |

**Table 2:** Results from the Linear Mixed Models run on data from the Adjust-Target and Move-To-Target tasks with the gain set as the dependent variable, with both Environment Structure and Ground Surface Condition set as fixed effects. This table reports differences in Environment Structure. This table reports unstandardized regression coefficients.

|  | Regression Coefficient | Standard Error | 95% CI (lower) | 95% CI (upper) | Significance |
| --- | --- | --- | --- | --- | --- |
| Corridor vs. Starfield (Move-To-Target Task) | 0.02 | 0.07 | -0.10 | 0.14 | n.s. |
| Corridor vs. Starfield (Adjust-Target Task) | 0.01 | 0.07 | -0.11 | 0.15 | n.s. |

#### Adjust-Target Task

The bottom row of Table 1 shows the means and standard deviations of the gains for the Adjust-Target task. Table 2 shows the results from the linear mixed model for the effect of Environment Structure on the gains of the Adjust-Target task. There were no significant differences between the corridor and starfield environment conditions. The gains are shown in Fig 3.

### Effect of the Ground Surface

#### Move-To-Target Task

The top row of Table 3 shows the means and standard deviations of the gains for the Move-To-Target task. Table 4 shows the results from the linear mixed model for the effect of the ground surface condition on the gains of the Move-To-Target task. We found that the grass texture led to significantly lower gains than the no texture or no ground surface conditions, these effects were small. We also found no significant differences between the no-texture and no-ground surface conditions. The gains are shown in Fig 3.

**Table 3:** Means and standard deviations of the gains for the different ground surface conditions.

|  | Grass Texture | No Texture | No Ground Surface |
| --- | --- | --- | --- |
| Move-To-Target Task | $1.19 \pm 0.51$ | $1.28 \pm 0.55$ | $1.31 \pm 0.62$ |
| Adjust-Target Task | $1.32 \pm 0.89$ | $1.20 \pm 0.72$ | $1.25 \pm 0.85$ |

**Table 4:** Results from the Linear Mixed Models run on data from the Move-To-Target and Adjust-Target tasks with the gain set as the dependent variable, with both Environment Structure and Ground Surface Condition set as fixed effects. This table reports differences in Environment Structure. This table reports unstandardized regression coefficients.

|  | Regression<br>Coefficient | Standard<br>Error | 95% CI<br>(lower) | 95% CI<br>(upper) | Significance |
| --- | --- | --- | --- | --- | --- |
| Grass Texture vs. No<br>Texture<br>(Move-To-Target Task) | 0.07 | 0.03 | 0.008 | 0.13 | * |
| Grass Texture vs. No<br>Ground Surface<br>(Move-To-Target Task) | 0.12 | 0.04 | 0.05 | 0.18 | * |
| No Texture vs. No<br>Ground Surface<br>(Move-To-Target Task) | -0.04 | 0.03 | -0.11 | 0.02 | n.s. |
| Grass Texture vs. No<br>Texture<br>(Adjust-Target Task) | -0.12 | 0.07 | 0.001 | 0.13 | * |
| Grass Texture vs. No<br>Ground Surface<br>(Adjust-Target Task) | -0.07 | 0.09 | 0.03 | 0.18 | * |
| No Texture vs. No<br>Ground Surface<br>(Adjust-Target Task) | -0.05 | 0.06 | -0.18 | 0.08 | n.s. |

#### Adjust-Target Task

The bottom row of Table 3 shows the means and standard deviations of the gains for the Adjust-Target task. Table 4 shows the results from the linear mixed model for the effect of ground surface condition on the gains of the Adjust-Target task. Contrary to the results from the Move-To-Target task, we found that in the Adjust-Target task, the grass texture led to significantly higher gains than the no texture or no ground surface conditions, although again, these effects were small. There were also no significant differences between no texture and no ground surface conditions. The gains are shown in Fig 3.

## Experiment 2: Naturalism

### Methods

#### Participants

Eighteen subjects (9F, 9M, mean age 20.2 yrs, SD ±2.8) participated in this study. The recruitment period was between January 23rd, 2024 and March 20th, 2024. All participants were recruited using the Kinesiology Undergraduate Participant Pool at York University. The protocols used in this study were approved by the York Human Participants Review Sub-committee (#e2021-407) and conducted in accordance with the Declaration of Helsinki. All participants gave prior informed written consent and were naive to the purpose of the study.

#### Apparatus

Same as Experiment 1.

#### Stimuli

There were three virtual street environments in which this experiment was performed (see Fig 4). The first condition was moving through a natural street scene where all the objects in the environment (people, trees, streetlights, houses) were scaled and oriented normally and remained stationary as the viewer moved past. The second “scaled up” condition, was a version of the virtual street where all the objects in the environment were twice as large as in the “natural” condition – if participants felt smaller or that distances were further apart but they still covered the same distances in the same time, then perceived speed might be expected to increase. The last condition was an unnatural condition in which all the objects in the environment were rotated and scaled randomly. In this last “unnatural” condition, each object was changed such that its original size was multiplied by a random number between 0.5 and 2.0, and rotated on each of its x, y, and z axes by a random number between +180 and −180 degrees, although in the y-axis the object was then set to the closest 0, 90, or 270 degree. The road and paving stones remained the same size and orientation in all conditions. The target (black and white checked pillar, measuring 1 x 5m) was presented in the same sparse black environment as Experiment 1 (see Fig 2).

**Fig 4.**
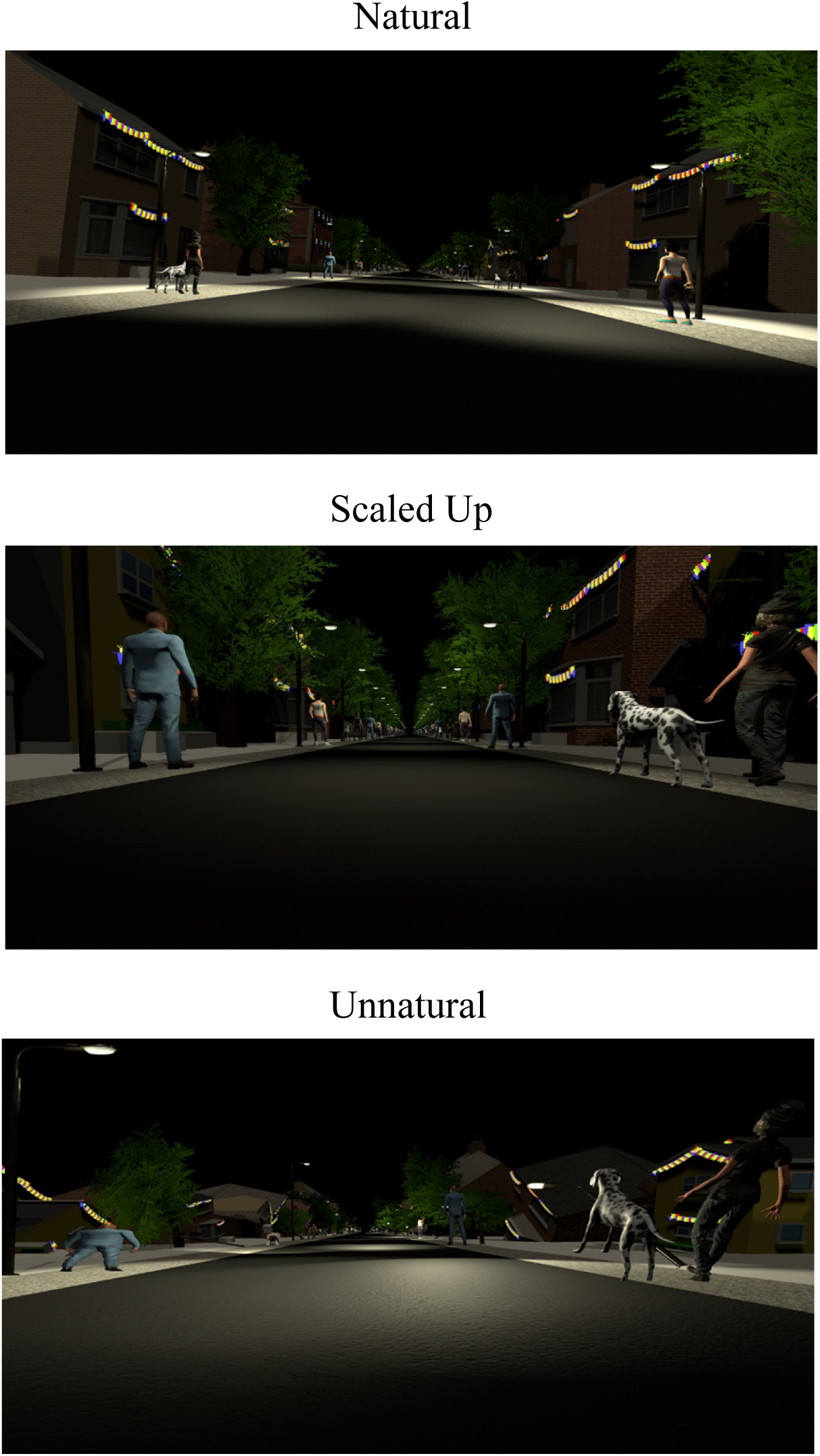
Visual scenes rendered inside the virtual reality headset. The top panel shows the natural condition, where objects in the environment were scaled normally. The central panel shows the scaled-up condition, where objects in the environment were 2x larger than the natural condition. The lower panel shows the unnatural condition, where objects in the environment were rotated and scaled randomly.

#### Procedure

Same as Experiment 1. Participants completed both the Move-To-Target and Adjust-Target tasks. Visual self-motion was simulated at 5 m/s down the street. Each task consisted of the same twelve target distances (5, 8, 11, 14, 17, 20, 23, 26, 29, 32, 35, 40 m), with no repetitions.

#### Data Analysis

Same as Experiment 1. Out of 1283 data points, 66 data points were removed using the same outlier analysis as Experiment 1. Using the same Barr et al.’s (2013) model comparison approach to determine the LMMs, random slopes for Environment per Participant were kept for both the Move-To-Target and Adjust-Target models. Therefore, the final model structure for the gains LMMs reads as follows:

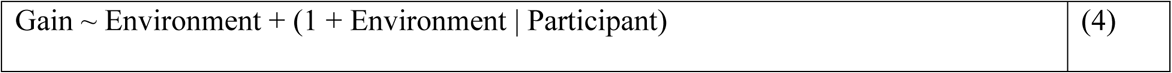

## Results

### Effect of Naturalism and Scale

#### Move-To-Target Task

The top row of Table 5 shows the means and standard deviations of the gains for the Move-To-Target task. Table 6 shows the results from the linear mixed model for the effect of Environment on the gains of the Move-To-Target task. We found that the scaled-up condition led to higher gains than the natural or unnatural conditions. However, we found no significant differences between the natural and unnatural conditions. The gains are shown in the top half of Fig 5.

**Fig 5.**
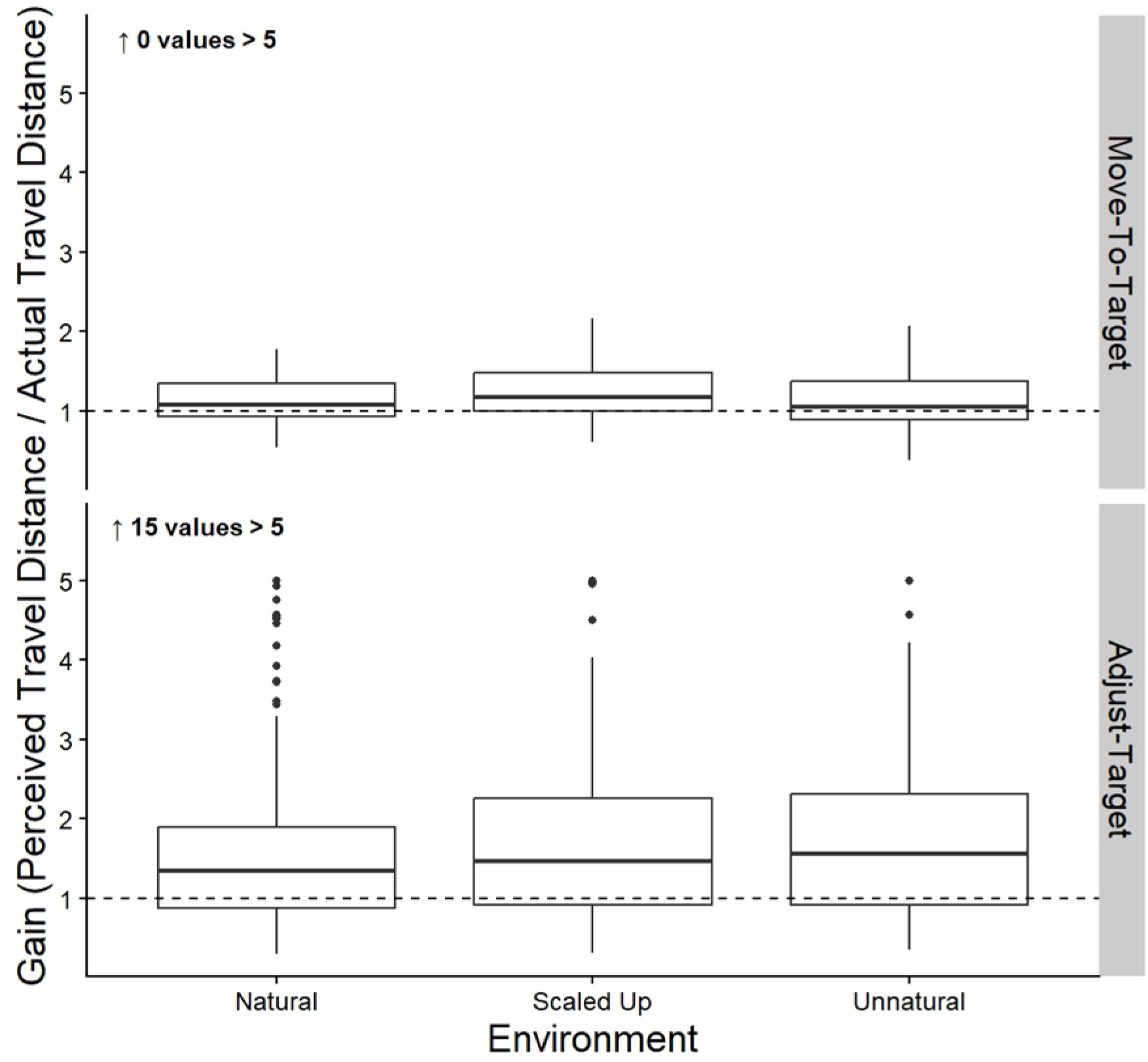
Gains. Box plots of the group gains for both the Move-To-Target (top row) and Adjust-Target (bottom row) tasks for each Environment. The middle line represents the median, the boxes extend from the first quartile to the third quartile, the whiskers extend up to 1.5 times the interquartile range, and the outliers are shown as individual points beyond the whiskers. The black dashed line represents perfect performance (gain of 1).

**Table 5:** Means and standard deviations of the gains for the different Environment conditions.

|  | Natural | Scaled Up | Unnatural |
| --- | --- | --- | --- |
| Move-To-Target Task | 1.12 ± 0.27 | 1.24 ± 0.33 | 1.12 ± 0.35 |
| Adjust-Target Task | 1.60 ± 1.02 | 1.82 ± 1.41 | 1.76 ± 1.12 |

**Table 6:** Results from the Linear Mixed Models run on data from the Move-To-Target and Adjust-Target tasks with the gain set as the dependent variable, with Environment Condition set as a fixed effect. This table reports differences in the Environment Condition. This table reports unstandardized regression coefficients.

|  | Regression Coefficient | Standard Error | 95% CI (lower) | 95% CI (upper) | Significance |
| --- | --- | --- | --- | --- | --- |
| Natural vs. Scaled Up (Move-To-Target Task) | 0.12 | 0.04 | 0.04 | 0.18 | * |
| Natural vs. Unnatural (Move-To-Target Task) | 0.008 | 0.05 | -0.08 | 0.11 | n.s. |
| Scaled Up vs. Unnatural (Move-To-Target Task) | -0.11 | 0.03 | -0.18 | -0.05 | * |
| Natural vs. Scaled Up (Adjust-Target Task) | -0.25 | 0.16 | -0.54 | 0.06 | n.s. |
| Natural vs. Unnatural<br>(Adjust-Target Task) | 0.18 | 0.08 | -0.08 | 0.11 | n.s. |
| Scaled Up vs. Unnatural<br>(Adjust-Target Task) | -0.08 | 0.16 | -0.37 | 0.23 | n.s. |

#### Adjust-Target Task

The bottom row of Table 5 shows the means and standard deviations of the gains for the Adjust-Target task. Table 6 shows the results from the linear mixed model for the effect of Environment on the gains of the Adjust-Target task. There were no significant differences between the natural, scaled up, or unnatural conditions. The gains are shown in the bottom half of Fig 5.

## Experiment 3: Colour

### Methods

#### Participants

Eighteen subjects (11F, 7M, mean age 19.9 yrs, SD ±2.8) participated in this study. The recruitment period was between September 28th, 2024 and November 3rd, 2024. All participants were recruited using the Kinesiology Undergraduate Participant Pool at York University. The protocols used in this study were approved by the York Human Participants Review Sub-committee (#e2021-407) and conducted in accordance with the Declaration of Helsinki. All participants gave prior informed written consent and were naive to the purpose of the study.

#### Apparatus

Due to experimental setup changes, a different virtual reality HMD and computer were used for this experiment. For Experiment 3, A VIVE Pro EYE HMD (field of view of 110°, resolution 1440 × 1600 per eye, 90 Hz refresh rate) was used to present the stimuli. The program was run on an Alienware laptop (16 GB RAM, Intel Core i7-9750H CPU, 2.60 GHz, NVIDIA GeForce RTX 2060).

#### Stimuli

There were two virtual environments in which this experiment was performed. Both environments were a starfield environment with a density of 3,000 stars. The size of the various “stars” in both starfield environments were the same as Experiment 1. The stars were randomly placed similar to Experiment 1, except in the y-axis, the stars were also placed below the horizon. The stars were placed anywhere between −25m and less than a random number between −2 or −5m, or between +25m and greater than a random number between +8 and 10m. The first environment was in colour, and the second was in greyscale (Fig 6). Both scenes had “stars” with various textures. Since luminance has been shown to influence vection (Guo et al., 2021), both scenes were matched in terms of luminance. The mean luminance of the two scenes was measured using a Colorimeter (Radiant I29). The colorimeter was positioned against the right lens of the headset. To keep the luminance levels the same (approximately 0.30), the global illumination of the grey scale environment within the Unity environment had to be reduced from 1.0 to 0.8. This value was calculated as the average from five different views in the headset (i.e., the program was restarted and measured five times, as the position of the spheres change on each trial). The target (black and white checked pillar, measuring 1 x 5m) was presented in the same sparse black environment as Experiments 1 and 2 (see Fig 2).

**Fig 6.**
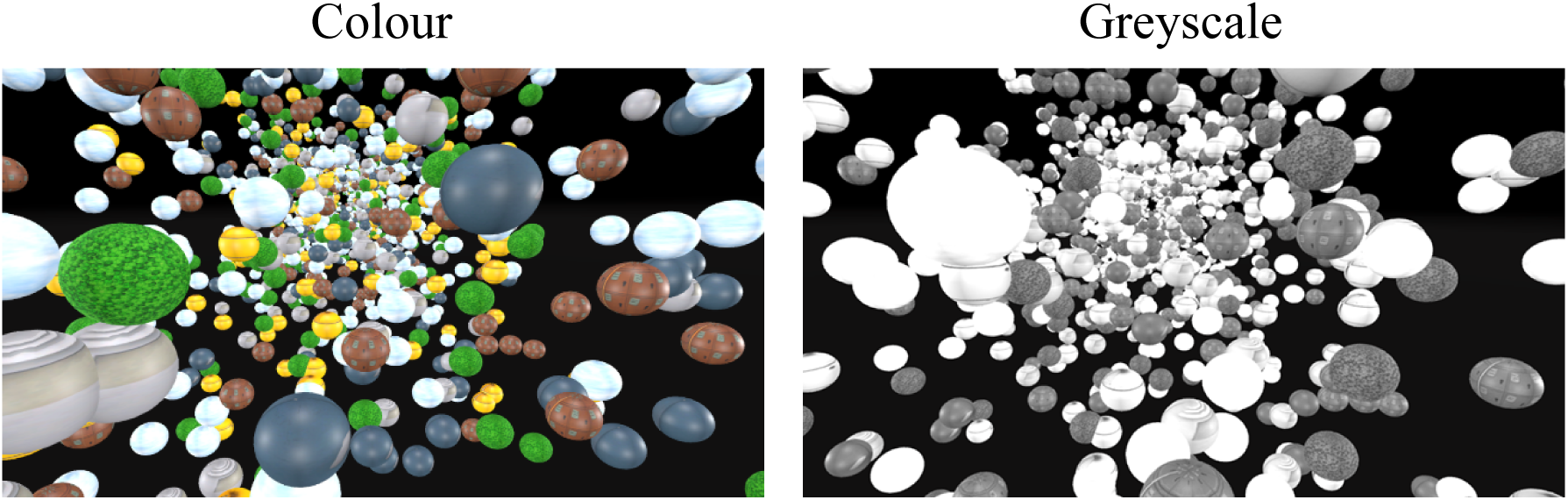
Visual scenes seen inside the virtual reality headset. The left panel shows the starfield of 3,000 stars in colour. The right panel shows the same starfield in greyscale.

#### Procedure

Same as Experiments 1 and 2.

#### Data Analysis

Out of 2,447 data points, 115 data points were removed using the same outlier analysis as Experiment 1. Like Experiment 2, random slopes for Environment per Participant were kept for both the Move-To-Target and Adjust-Target LMM models, therefore we used the same model structure for both LMMs:

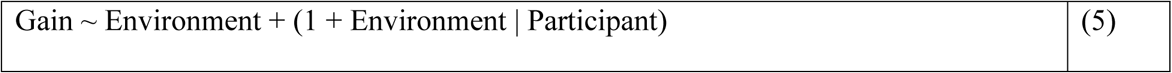

## Results

### Effect of Colour

#### Move-To-Target Task

The top row of Table 7 shows the means and standard deviations of the gains for the Move-To-Target task. Table 8 shows the results from the linear mixed model for the effect of Colour on the gains of the Move-To-Target task. We found no significant differences between the colour and greyscale conditions. The gains are shown in the top half of Fig 7.

**Fig 7.**
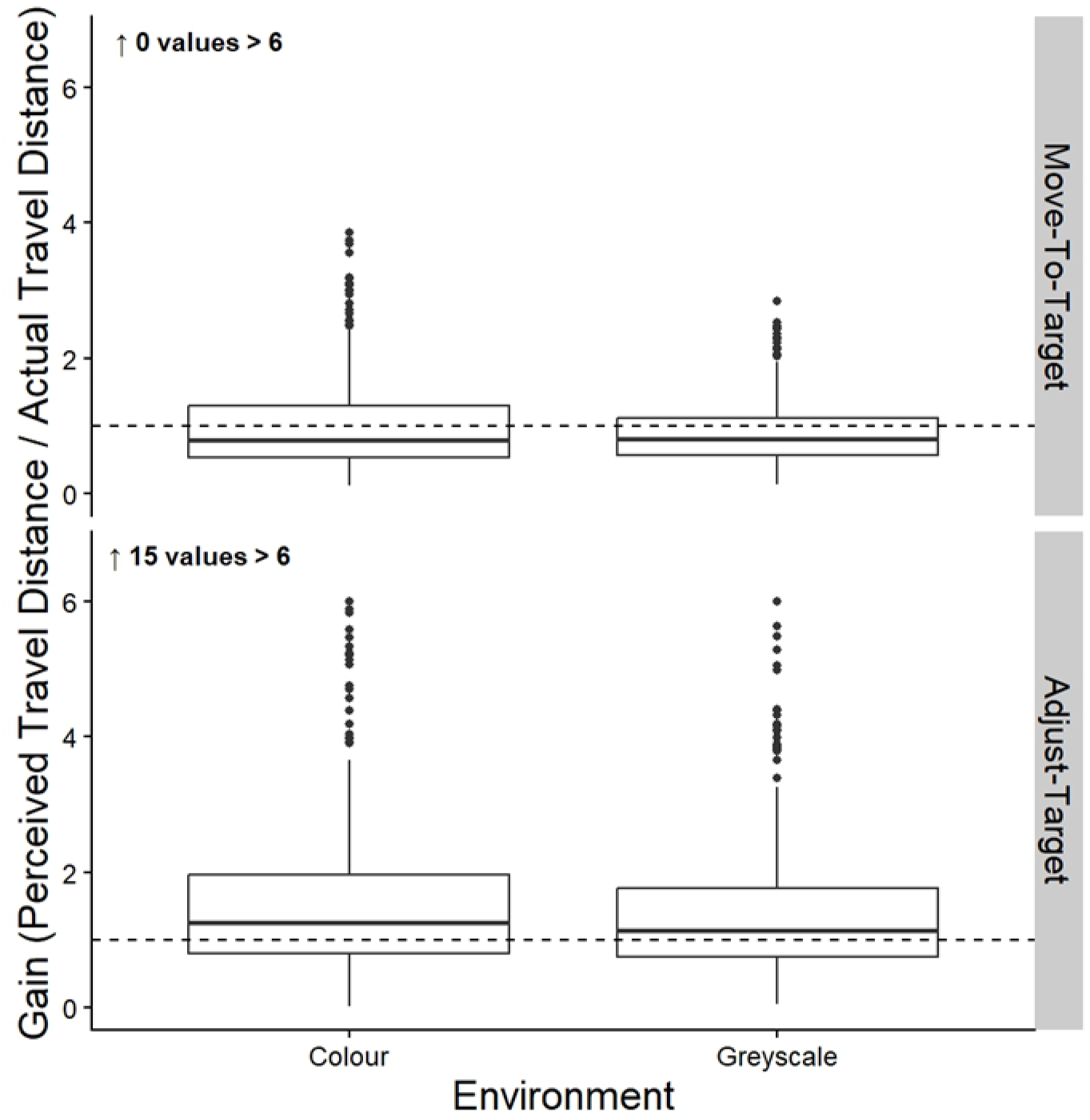
Gains. Box plots of the group gains for both the Move-To-Target (top row) and Adjust-Target (bottom row) tasks for the Colour and Greyscale conditions. The middle line represents the median, the boxes extend from the first quartile to the third quartile, the whiskers extend up to 1.5 times the interquartile range, and the outliers are shown as individual points beyond the whiskers.

**Table 7:**
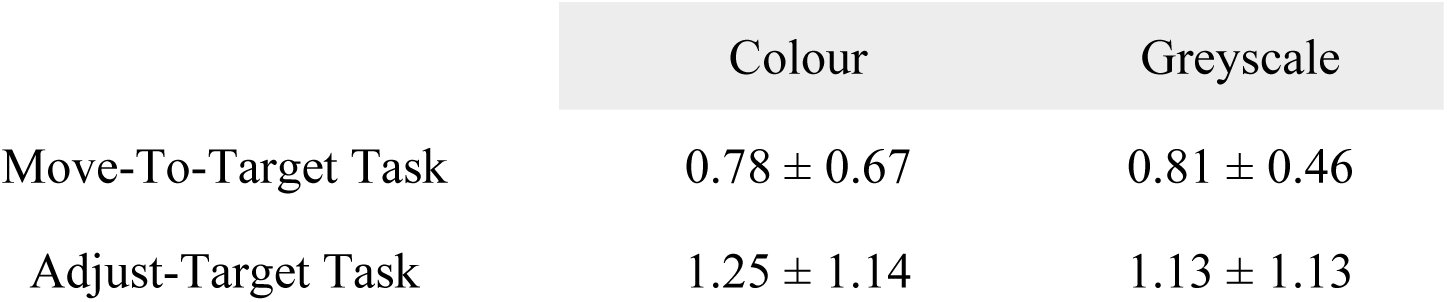
Means and standard deviations of the gains for the coloured and greyscale conditions.

|  | Colour | Greyscale |
| --- | --- | --- |
| Move-To-Target Task | $0.78 \pm 0.67$ | $0.81 \pm 0.46$ |
| Adjust-Target Task | $1.25 \pm 1.14$ | $1.13 \pm 1.13$ |

**Table 8:**
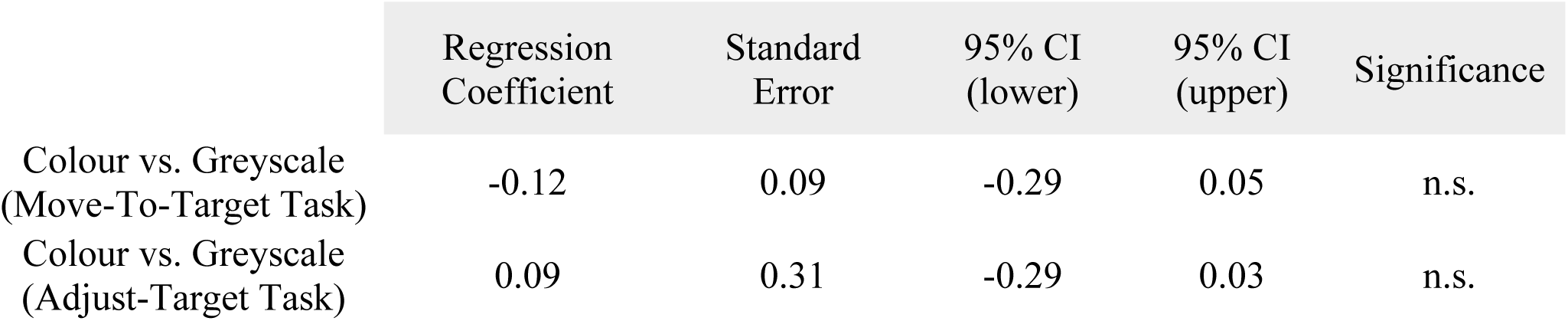
Results from the Linear Mixed Models run on data from the Move-To-Target and Adjust-Target tasks with the gain set as the dependent variable, with Colour Condition set as the fixed effect. This table reports differences in the Colour. This table reports unstandardized regression coefficients.

#### Adjust-Target Task

The bottom row of Table 7 shows the means and standard deviations of the gains for the Adjust-Target task. Table 8 shows the results from the linear mixed model for the effect of Colour on the gains of the Adjust-Target task. There were no significant differences between the colour and greyscale conditions. The gains are shown in the bottom half of Fig 7.

## Experiment 4: Starfield Density

### Methods

#### Participants

Eighteen subjects (11F, 7M, mean age 19.5 yrs, SD ±2.1) participated in this study. The recruitment period was between February 15th, 2025 and Apil 11th, 2025. All participants were recruited using the Kinesiology Undergraduate Participant Pool at York University. The protocols used in this study were approved by the York Human Participants Review Sub-committee (#e2021-407) and conducted in accordance with the Declaration of Helsinki. All participants gave prior informed written consent and were naive to the purpose of the study.

#### Apparatus

Same as Experiment 3.

#### Stimuli

In this experiment, we were interested in the starfield density needed to accurately estimate travel distance. To do this, we manipulated the density of stars in the starfield environment to either 10 stars, 100 stars, or 1,000 stars in the environment at a time (Fig 8), as well as moving participants at three different travel speeds (1, 5, 10 m/s). The stars’ locations were set to the same parameters as Experiment 3 and had a white and faint blue texture. The target (black and white checked pillar, measuring 1 x 5m) was presented in the same sparse black environment as Experiments 1-3 (see Fig 2).

**Fig 8.**
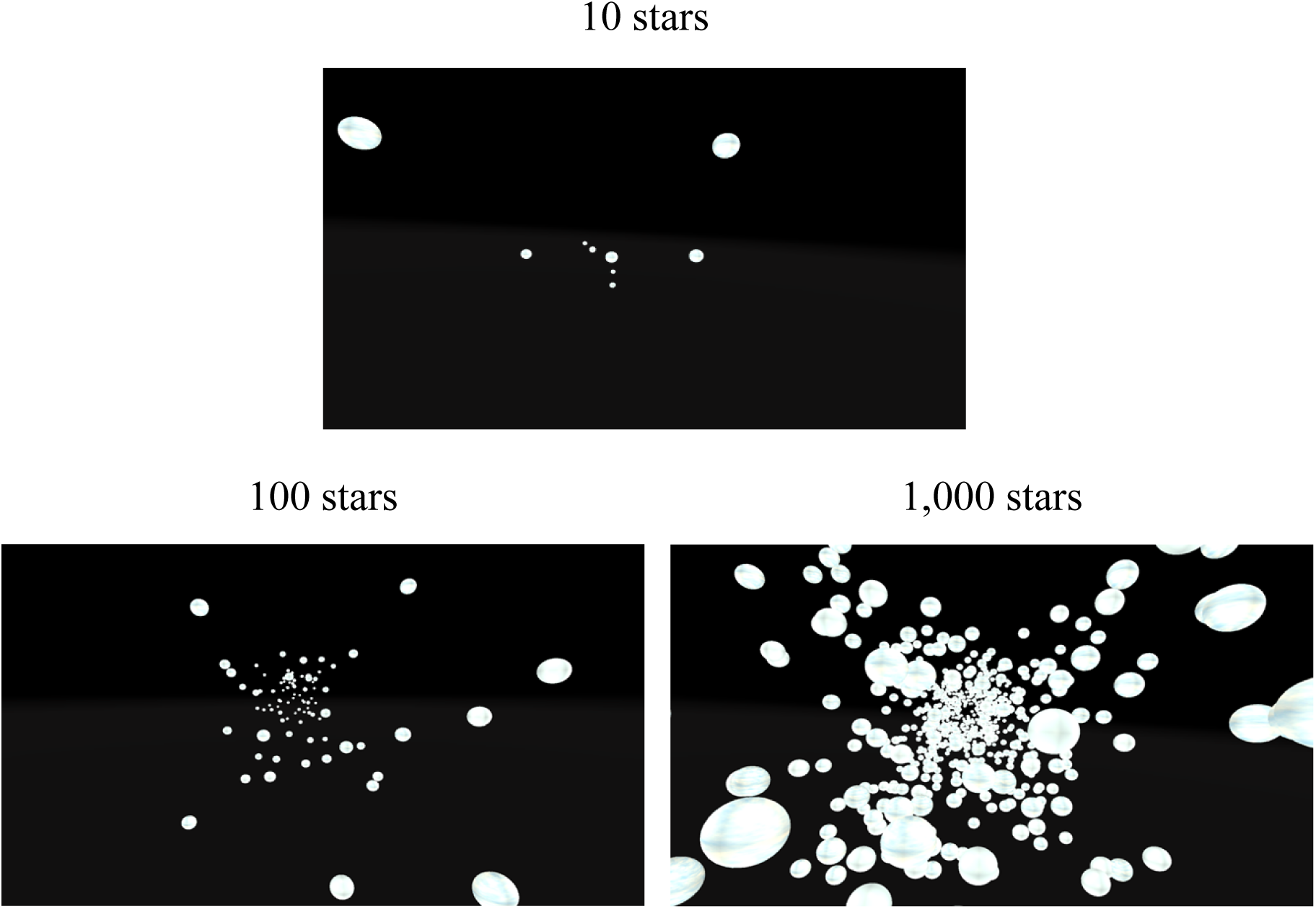
Visual scenes seen inside the virtual reality headset. The top panel shows the low-density condition, which contained 10 stars. The bottom left panel shows the medium-density condition, which contained 100 stars. The bottom right panel shows the high-density condition, which contained 1,000 stars.

#### Procedure

The procedure was the same as Experiments 1-3, with minor changes. Self-motion was simulated at three different speeds (1, 5, 10 m/s), the order of which was randomized in each block. Each block consisted of six target distances (10, 15, 20, 25, 30, 40 m). As in Experiments 1-3, these two tasks were counterbalanced such that half of the participants completed the Move-To-Target first and half completed the Adjust-Target task first.

#### Data Analysis

Data analysis used the same methods as used in Experiment 1. Out of 1,152 data points, 95 data points were removed using the same outlier analysis mentioned above. For the Linear Mixed Model, the most appropriate model structure was determined using Barr et al.’s (2013) model comparison approach. We started with a maximal model including all relevant experimental variables (Starfield Density and Speed) as slopes per participant and compared to simpler models until no significant differences were found between models. Only random slopes for Speed per Participant were kept for both the Move-To-Target and Adjust-Target models. We did not include an interaction between Starfield Density and Speed because we had no specific hypotheses about the interaction term. The final model structure for the LMM on the gains reads as follows:

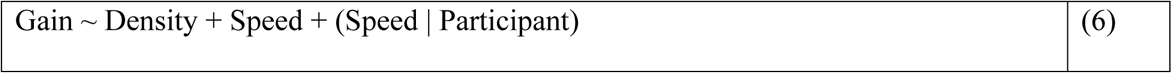

## Results

### Effect of Starfield Density

#### Move-To-Target Task

The top row of Table 9 shows the means and standard deviations of the gains for the Move-To-Target task. Table 10 shows the results from the linear mixed model for the effect of Starfield Density on the gains of the Move-To-Target task. We found a significant difference between the 10- and 100-star conditions, although there were no significant differences between the 10- and 1,000-starfield-density conditions or the 100- and 1,000-starfield-density conditions. The gains are shown at the top of Fig 9.

**Fig 9.**
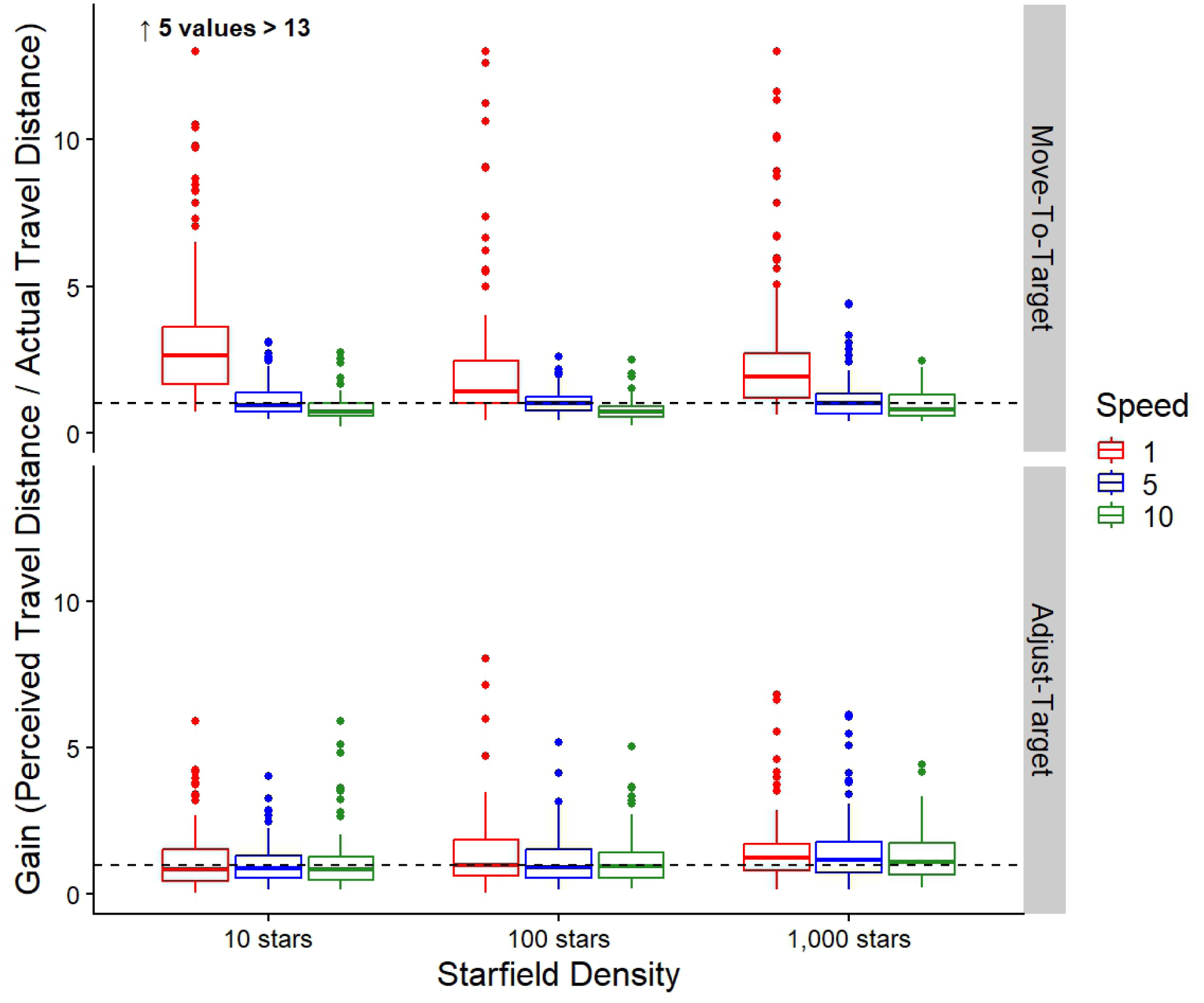
Gains. Box plots of the group gains for both the Move-To-Target (top row) and Adjust-Target (bottom row) tasks for each Starfield Density and Speed Condition. The middle line represents the median, the boxes extend from the first quartile to the third quartile, the whiskers extend up to 1.5 times the interquartile range, and the outliers are shown as individual points beyond the whiskers. The 1 m/s is represented in red (left most), 5 m/s in blue (second from left), and 10 m/s in green (right most). The black dashed line represents perfect performance (gain of 1).

**Table 9:** Means and standard deviations of the gains for the Starfield Density conditions.

|  | 10 stars | 100 stars | 1,000 stars |
| --- | --- | --- | --- |
| Move-To-Target Task | 2.15 ± 2.92 | 1.68 ± 2.57 | 2.24 ± 2.84 |
| Adjust-Target Task | 1.27 ± 1.07 | 1.40 ± 1.25 | 1.67 ± 1.29 |

**Table 10:** Results from the Linear Mixed Models run on data from the Move-To-Target and Adjust-Target tasks with the gain set as the dependent variable, with both Starfield Density and Speed set as fixed effects. This table reports differences in Starfield Density. This table reports unstandardized regression coefficients.

|  | Regression Coefficient | Standard Error | 95% CI (lower) | 95% CI (upper) | Significance |
| --- | --- | --- | --- | --- | --- |
| 10 stars vs. 100 stars (Move-To-Target Task) | 0.84 | 0.39 | 0.16 | 1.61 | * |
| 10 stars vs. 1,000 stars (Move-To-Target Task) | 0.19 | 0.39 | -0.54 | 1.02 | n.s. |
| 100 stars vs. 1,000 stars (Move-To-Target Task) | -0.65 | 0.39 | -1.39 | 0.17 | n.s. |
| 10 stars vs. 100 stars<br>(Adjust-Target Task) | 0.11 | 0.15 | -0.22 | 0.44 | n.s. |
| 10 stars vs. 1,000 stars<br>(Adjust-Target Task) | -0.42 | 0.15 | -0.60 | 0.92 | n.s. |
| 100 stars vs. 1,000 stars<br>(Adjust-Target Task) | -0.30 | 0.15 | -1.53 | 0.19 | n.s. |

#### Adjust-Target Task

The bottom row of Table 9 shows the means and standard deviations of the gains for the Adjust-Target task. Table 10 shows the results from the linear mixed model for the effect of Starfield Density on the gains of the Adjust-Target task. We found no significant differences between the 10, 100, and 1,000 starfield density conditions. The gains are shown at the bottom of Fig 9.

### Effect of Speed

#### Move-To-Target Task

The top row of Table 11 shows the means and standard deviations of the gains for the Move-To-Target task. Table 12 shows the results from the linear mixed model for the effect of Speed on the gains of the Move-To-Target task. We found that moving at 1 m/s led to significantly higher gains than the 5 m/s or 10 m/s. However, we found no significant differences between the 5 m/s and 10 m/s speed conditions. The gains are shown in Fig 9.

**Table 11:**
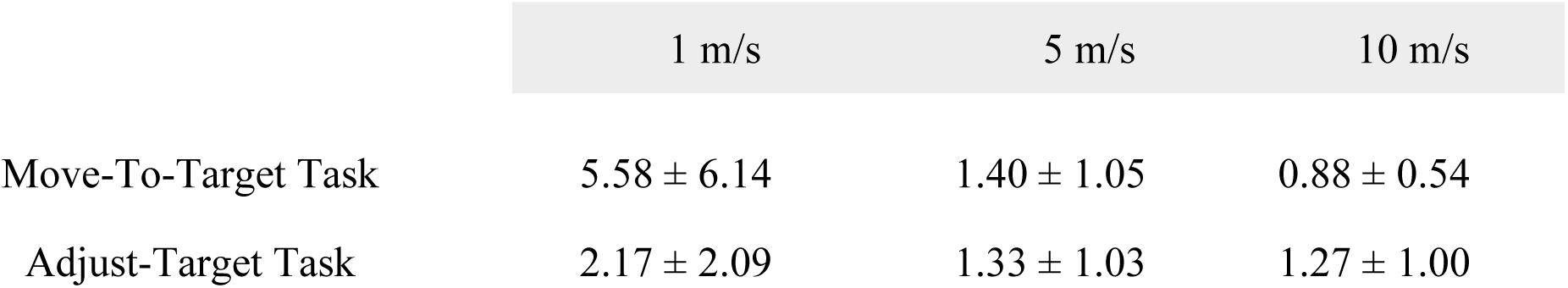
Means and standard deviations of the gains for the different speed conditions averaged across the 10-, 100-, and 1,000-star conditions.

**Table 12:** Results from the Linear Mixed Models run on data from the Move-To-Target and Adjust-Target tasks with the gain set as the dependent variable, with both Starfield Density and Speed set as fixed effects. This table reports differences in Starfield Density. This table reports unstandardized regression coefficients.

|  | Regression Coefficient | Standard Error | 95% CI (lower) | 95% CI (upper) | Significance |
| --- | --- | --- | --- | --- | --- |
| 1 m/s vs 5 m/s (Move-To-Target Task) | -4.20 | 1.15 | -6.26 | -2.19 | * |
| 1 m/s vs. 10 m/s (Move-To-Target Task) | -4.70 | 1.23 | -7.08 | -2.43 | * |
| 5 m/s vs. 10 m/s (Move-To-Target Task) | 0.50 | 0.35 | -0.28 | 1.22 | n.s. |
| 1 m/s vs 5 m/s (Adjust-Target Task) | -0.89 | 0.25 | -6.43 | -1.83 | * |
| 1 m/s vs. 10 m/s (Adjust-Target Task) | -0.94 | 0.26 | -7.15 | -2.33 | * |
| 5 m/s vs. 10 m/s (Adjust-Target Task) | 0.05 | 0.14 | -0.22 | 0.35 | n.s. |

#### Adjust-Target Task

The bottom row of Table 11 shows the means and standard deviations of the gains for the Adjust-Target task. Table 12 shows the results from the linear mixed model for the effect of Speed on the gains of the Adjust-Target task. Similar to the Move-To-Target task, we found that the 1 m/s condition led to significantly higher gains than the 5 m/s and 10 m/s conditions. However, there were no significant differences between the 5 and 10 m/s speed conditions. The gains are shown in Fig 9.

## Discussion

This study comprised a series of four experiments investigating how the characteristics of a virtual environment might affect the perception of travel distance. Aligning with our original hypotheses, we found that the texture of the ground surface (Experiment 1) and the scale of the environment (Experiment 2) had an effect on the perception of travel distance, although, these effects were small and should not be overemphasized. Contrary to our hypotheses, however, we found that manipulating the structure, the presence or absence of a ground surface (Experiment 1), the naturalism (Experiment 2), and the presence of colour (Experiment 3) in the visible scene had no effect on the ability of the observer to extract their movement information correctly and generate a consistent perception of distance moved. Experiment 4 confirmed the presence of a ceiling effect in the amount of stars needed to provide adequate optic flow to accurately estimate travel distance.

### Experiments 1-3: The Effect of Structure, Ground Surface, Naturalism, Scale, Colour

Much of the research on self-motion perception has used vection as a measurement. Although the different measures of the experience of vection can inform us as to whether people feel like they are moving, one can still perceive travel distance without experiencing vection. This divergence aligns with previous research that shows an increase in vection speed in microgravity (Oman et al., 2003; Young & Shelhamer, 1990) but no differences in perceived travel distance in microgravity when it is actually measured (Jörges et al., 2024). Others have also found an effect of the speed of optic flow on vection speed (De Graaf et al., 1990) but there is still mixed evidence as to whether there is an effect of speed on perceived travel distance (Bansal et al., 2024; Frenz et al., 2007; Harris et al., 2012; Lappe et al., 2007; McManus et al., 2017). Although McManus and Fiehler (2025) have shown that the perception of travel distance and the sensation of vection should be related, the studies quoted above highlight the idea that the sensation of vection might be independent of the perception of travel distance. There are two ways in which you can perceive self-motion without experiencing the sensation of vection: (1) you can experience that you are moving through a virtual environment without feeling like you are actually moving (e.g., when playing a first-person video game), (2) optic flow information can be interpreted not just as the sensation that you are moving through the world (i.e., vection) but also as you staying stationary while the world is moving past you. Many studies that have tested how the components of a virtual environment can affect your perceived self-motion have focused on the experience of vection (Bubka & Bonato, 2010; Seno et al., 2010; Tamada & Seno, 2015; Vaziri-Pashkam & Cavanagh, 2008), but no one has previously tested how the components of a virtual environment affect the perception of travel distance specifically. Our findings from Experiments 1-3 show no significant differences in travel distance estimates when manipulating the structure, naturalism, or colour of a virtual environment. In Experiment 2, we did find minor differences in scale, where the scaled-up condition led to higher gains in the Move-To-Target task than the natural or unnatural conditions. Although no significant differences were found in the Adjust-Target task. One limitation to Experiment 2 on naturalism, is that in all conditions, the roads, sidewalks, and participant eye height to the ground remained the same. In the unnatural condition, the road and sidewalk could have given natural cues. In the scaled-up condition, the participant eye height to the ground could have given a cue for scale. In Experiment 1, we did find significant differences related to the texture of the ground surface, although these effects were small. We find the textured ground led to lower gains in the Move-To-Target and higher gains in the Adjust-Target task. Again, since these effects were small, we do not want to overemphasize these findings. Generally, we show that the differences previously reported in vection do not necessarily translate to differences in perceived travel distance.

### Experiment 4 – The Effect of Starfield Density

Experiment 4 was conducted with the hope of shedding light on why we did not find differences in perceived travel distance between different virtual environments. We thought that our manipulations over the first three experiments were perhaps not strong enough to overcome the fact that there was still enough optic flow information to accurately make these judgements. In Experiment 4, we were interested in whether there may be a ceiling effect on the ability to extract the amount of optic flow needed to accurately estimate travel distance and that the reason we found no effect with these other manipulations was because that threshold had already been exceeded. To test this, we manipulated the density of stars in a starfield environment. Our results showed that travel distance could still be accurately estimated even with very little optic flow information, meaning that people might be very sensitive to optic flow information leading to a ceiling effect with how much optic flow is needed to accurately perceive travel distance. We hypothesize that this is why we did not find significant differences between many of the conditions in our first three experiments. The minor manipulations we made between structure, naturalism, and colour conditions did not integrally alter the environment enough to modulate the perception of travel distance because there was always enough optic flow information to make accurate judgements of travel distance, though we do not know exactly what the perceptual threshold for the amount of optic flow needed for travel distance estimation. In a future study, it would be interesting to add other movement cues e.g., a starfield in which some percentage of stars/dots obey the optic flow rules and the others are moving in random direction, to obtain coherence thresholds of the system that calculates distance travelled from optic flow to fully understand how much optic flow is needed to accurately perceive travel distance, similar to Burr and Santoro (2001) who calculated the perceptual threshold needed to discern the direction of motion of a random dot pattern.

### Conclusions

Here, we show that many of the characteristics of a virtual scene have no effect on the perception of travel distance within the limits of how we varied them. This line of research provides insights into the effects of a ground surface on perceived travel distance, which will help in designing helicopter and airplane simulators for training pilots to fly using the optic flow information provided by the ground surface. Together this series of experiments has implications for the design of virtual environments where estimating travel distance is important.

